# GRAMMAR-Lambda: An Extreme Simplification for Genome-wide Mixed Model Association Analysis

**DOI:** 10.1101/2021.03.10.434574

**Authors:** Runqing Yang, Jin Gao, Yuxin Song, Zhiyu Hao, Pao Xu

**Author notes:** **Corresponding Authors**: Runqing Yang, Pao Xu.

## Abstract

A highly efficient genome-wide association method, GRAMMAR-Lambda is proposed to make simple genomic control for the test statistics deflated by GRAMMAR, producing statistical power as high as exact mixed model association method. Using the simulated and real phenotypes, we show that at a moderate or above genomic heritability, polygenic effects can be estimated using a small number of randomly selected markers, which extremely simplify genome-wide association analysis with an approximate computational complexity to naïve method in large-scale complex population. Upon a test at once, joint association analysis offers significant increase in statistical power over existing methods.

## Introduction

Population stratification, family structure, and cryptic relativeness, as widespread confounding factors, inflate test statistics in genome-wide association studies (GWAS), which increases the false-positive rate of association tests. The extent of this inflation is measured using genomic control ^1^. The most straightforward approach to correct these confounding effects is to divide genome-wide test statistics by genomic control calculated as the average or median genome-wide test statistics ^2–4^. Principal component analysis and family-based association tests are applied to correct population stratification ^5–10^ and family structure ^11–14^, respectively. In comparison, the linear mixed-model (LMM) ^15,16^ considers not only population stratification and family structure, but also cryptic relatedness through known kinship, offering a practical and comprehensive genomic control approach.

Even though the genomic relationship matrix (GRM) ^17^, calculated with whole genomic markers, is used to estimate corresponding polygenic variances to candidate markers, the LMMs are much more computation-intensive for high-throughput single nucleotide polymorphisms (SNPs). Several simplified algorithms for mixed model association have been developed to improve computational efficiency. Assuming that a quantitative trait nucleotide (QTN) has a minor variance, both EMMAX ^18^ and P3D ^19^ replaced different polygenic variances with the same genomic variance estimated by null LMM. Subsequently, FaST-LMM ^20^ and GEMMA ^21^ use a single eigen decomposition of the GRM ^22^ to reduce the estimation of polygenic or genomic variances. After moving the polygenic effects estimated by the pedigree-based best linear unbiased prediction (PBLUP) ^15^ from phenotypes, initial GRAMMAR ^23^ associates each SNP with phenotype residuals. Instead of the PBLUP, however, GRM-based best linear unbiased prediction (GBLUP) ^24,25^ increases GRAMMAR’s false-negative error rate because of accurately estimating genomic breeding values (GBVs) ^17,24,25^. Therefore, GRAMMAR-Gamma ^26^ and BOLT-LMM ^27^ have been proposed to improve the statistical power of QTN detection by calibrating the deflated test statistics by the GRAMMAR.

Before association tests, all LMM-based methods are required to calculate GRM and estimate variance components. Using these estimated components, GRAMMAR determines the least computing time for association tests, *O*(*nm*), where *n* is the population size and *m* is the number of markers. Based on the GRAMMAR, BOLT-LMM estimates variance components with Monte Carlo REML ^28,29^ only in the computing complexity of *O*(*inm*) ^27^, where *i* is iterative times. For large-scale data of humans, a resource-efficient tool for mixed-model association analysis has been developed to enhance computing efficiency of GRAMMAR-Gamma significantly, via calculating sparse GRM ^30^.

In this study, we propose a GRAMMAR-Lambda method to adjust GRAMMAR using genomic control, extremely simplifying genome-wide mixed model analysis. We show, using extensive simulations, that for a complex population structure, a high false-negative error of GRAMMAR can be efficiently corrected by dividing genome-wide test statistics by genomic control. Statistical power is kept constant, almost the same as that of exact FaST-LMM, even with randomly selected markers across the whole genome. In estimating GBVs with GBLUP, variance components or genomic heritability do not require to be estimated in advance. Within the framework of the GRAMMAR-Lambda, the joint analysis for QTN candidates significantly improves the statistical power, as compared to a test at once.

## Results

### Statistical property of GRAMMAR-Lambda

GRAMMAR-Lambda, a test at once was comparable to the four competing methods, such as FaST-LMM, GRAMMAR, GRAMMAR-Gamma and BOLT-LMM. Association results from the simulation (1) are displayed selectively in Figure 1 for Q-Q profiles and Figure 2 for ROC profiles, and genomic control values are calculated in Table 1S. In general, GRAMMAR-Lambda showed stable and good statistical properties that did not depend on how many the simulated QTNs were responsible for quantitative traits and how complex the population structures were. Among GRAMMAR-Lambda and four competing methods, GRAMMAR had the lowest genomic controls and statistical powers. At the same time, it showed more complex population structures and more strong deflation of genome-wide test statistics. Under perfect genomic control of exact 1, GRAMMAR-Lambda could detect QTNs with the slightly high statistical powers, compared to FaST-LMM and GRAMMAR-Gamma. Although GRAMMAR-Gamma and BOLT-LMM achieved genomic control for GRAMMAR via different calibration factors 26,27, GRAMMAR-Gamma a little deflates test statistics for complex maize population, making the genomic control below 1.0. On the contrary, BOLT-LMM inflates test statistics, so that increases the false-positive rate as the complexity of population structure increased.

**Figure 1:**
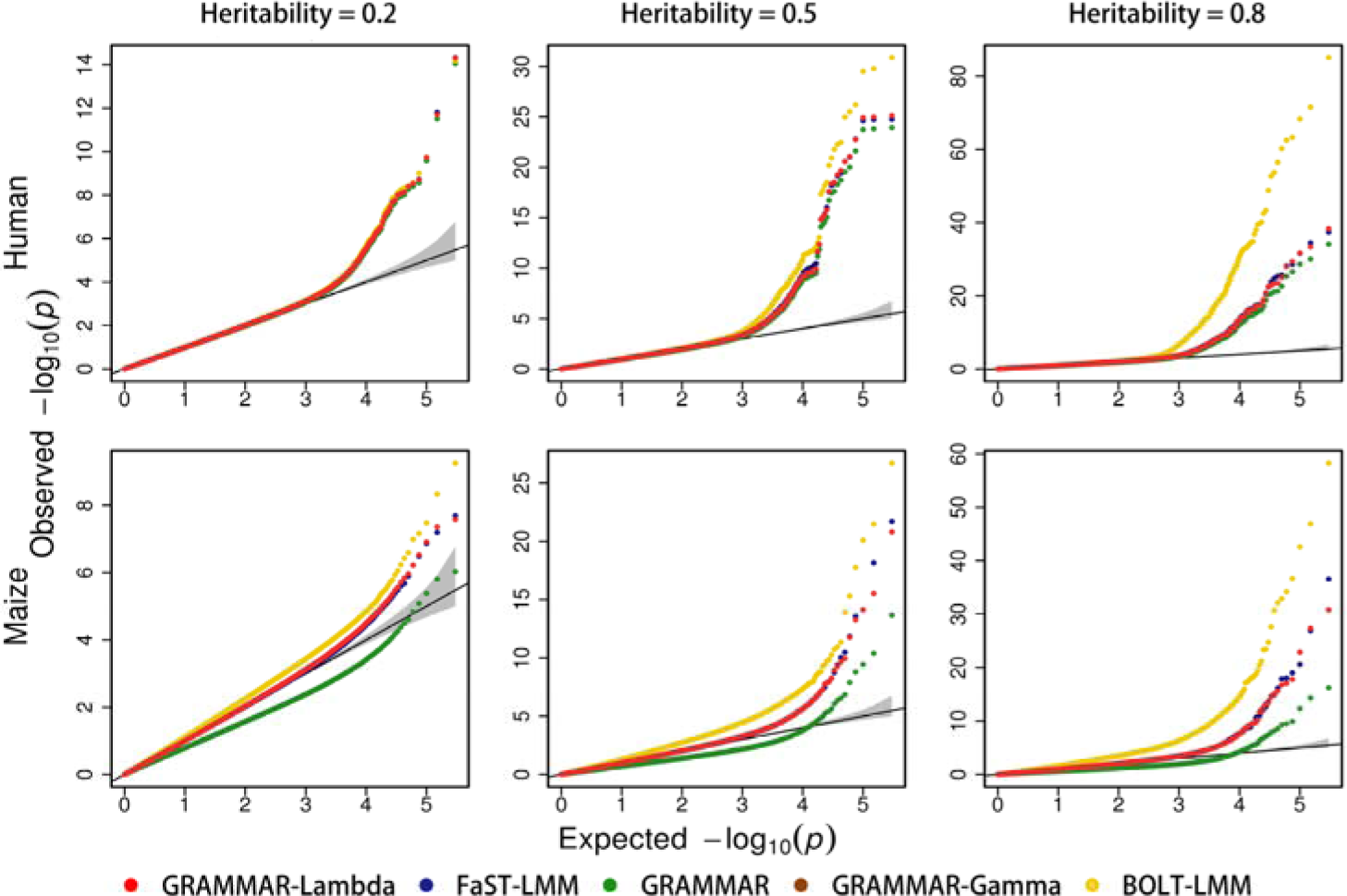
Comparison in the Q-Q plots between GRAMMAR-Lambda and the four competing methods. The simulated phenotypes are controlled by 200 QTNs with the low, moderate and high heritabilities in human and maize. The Q-Q plots for all simulated phenotypes are reported in Supplementary Figure 1S.

**Figure 2:**
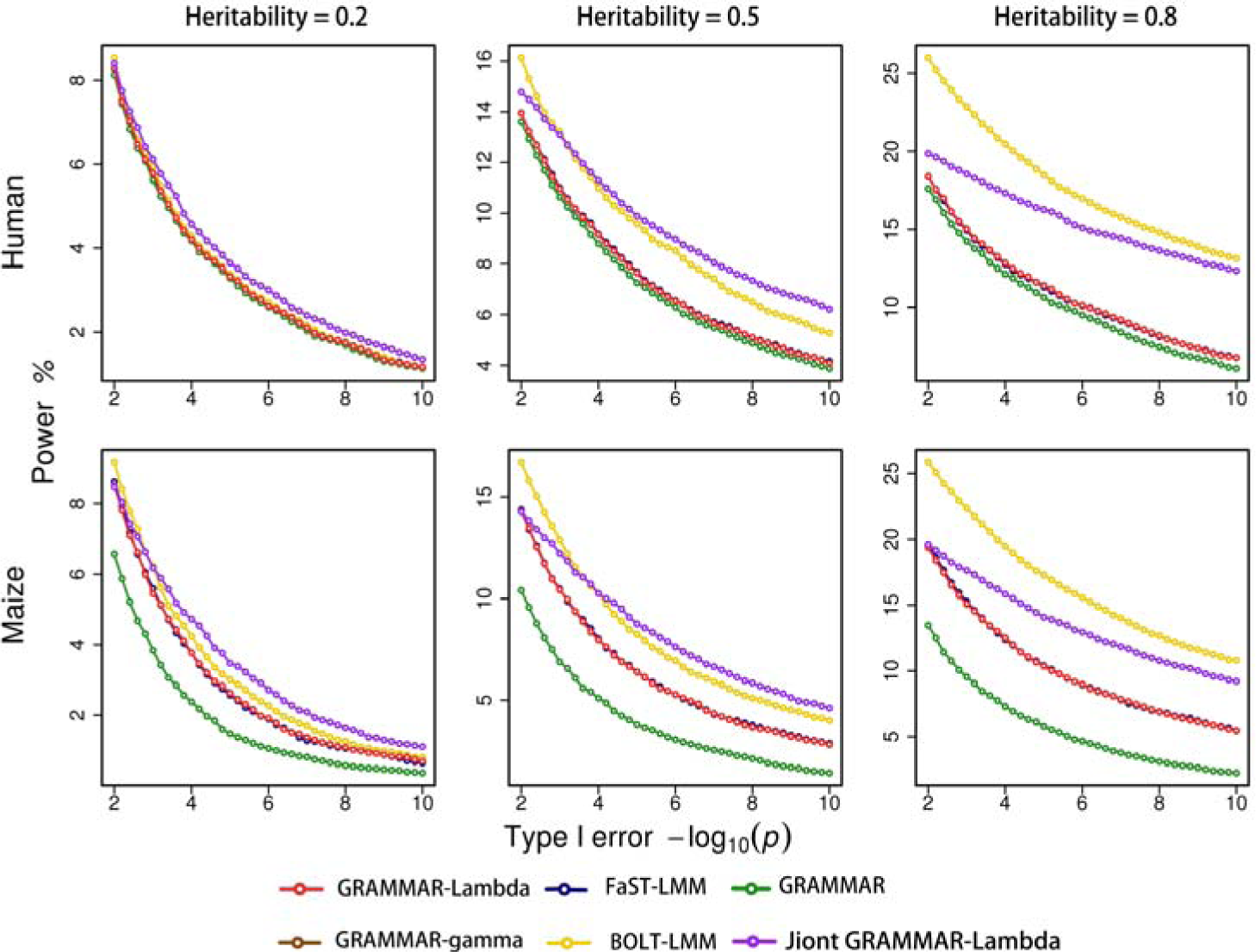
Comparison in the ROC profiles between GRAMMAR-Lambda and the four competing methods. The ROC profiles are plotted using the statistical powers to detect QTNs relative to the given series of Type I errors. Here, the simulated phenotypes are controlled by 200 QTNs with the low, moderate and high heritabilities in human and maize. The ROC profiles for all simulated phenotypes are reported in Supplementary Figure 2S.

After genomic control for GRAMMAR, GRAMMAR-Lambda jointly analysed multiple QTN candidates chosen with a test at once, at a significance level of 0.05. For convenience to compare, the statistical powers with joint association analyses were depicted along with those using a test at once. Though backward multiple regression analysis, GRAMMAR-Lambda gained significant increase in statistical powers. Although BOLT-LMM also showed an increase in the statistical power to detect QTNs, it did not control the false-positive error, especially for the complex maize population.

### Sensitivity to estimate genomic heritability

All LMM-based GWASs, except for GRAMMAR, required the exact estimation of polygenic variances or effects. Since GRAMMAR-Lambda efficiently controlled the deflation of test statistics caused by overestimation of polygenic effects, we hypothesised that with GBLUP, the appropriate estimation or overestimation of polygenic effects could be produced by pre-setting a genomic heritability value greater than the actual genomic heritability or no less than 50% of it. Note that the given lower bound of genomic heritability was obtained via our extensive simulations for optimising GRAMMAR (not shown).

With the specified genomic heritabilities of 0.05, 0.1, 0.2, 0.3, 0.5, 0.7, 0.8, 0.9 and 0.95, we analysed the phenotypes from the simulation (2) using only GRAMMAR-Lambda to examine the sensitivity to estimate genomic heritability. Figure 3 compared statistical powers generated from the estimated genomic heritabilities with those from the specified genomic heritabilities. As shown in each plot for human population, most ROC curves of the estimated genomic heritabilities almost overlapped with those of pre-specified genomic heritabilities. The maximum thickness of each overlapped curve was less than one QTN converted by the difference in the statistical power. For the simulated phenotypes in maize, however, the decrease in statistical powers was observed at very low or high genomic heritabilities simulated. Despite this, GRAMMAR-Lambda could retain high statistical powers with perfect genomic controls (see Figure 3S), when genomic heritability was chosen from any value within a broader interval to estimate GBVs. Thus, there was no need to estimate variance components or genomic heritability. When implementing GRAMMAR-Lambda, genomic heritability was set to 0.5 by default or empirical heritability of traits if available.

**Figure 3:**
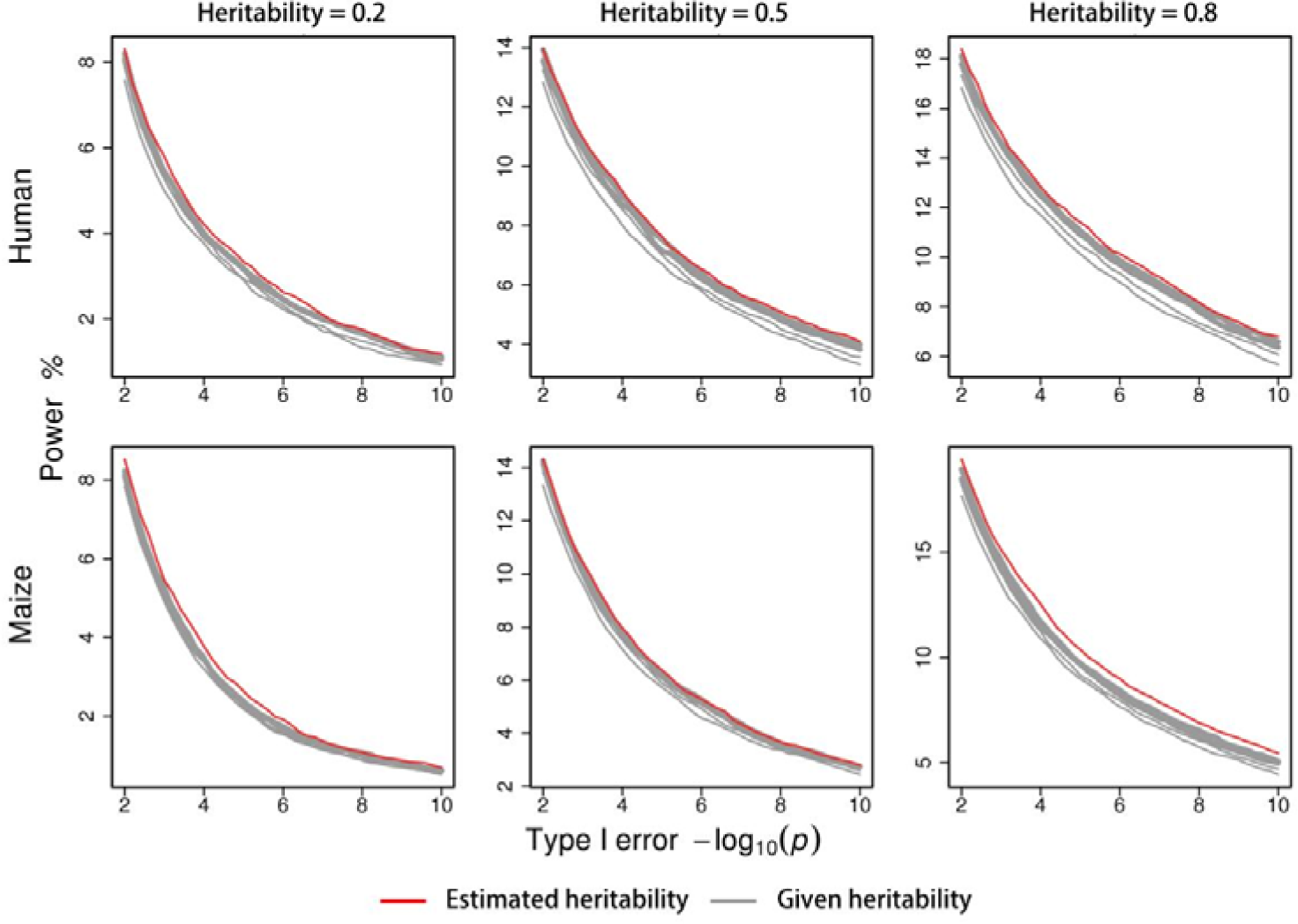
Sensitivity of statistical powers to the specified heritabilities for GRAMMAR-Lambda. Statistical powers are dynamically evaluated with the ROC profiles. The simulated phenotypes are controlled by 200 QTNs with the low, moderate and high heritabilities in human and maize.

### Calculation of GRMs with the sampling markers

Estimation of genomic heritability and GBVs using GBLUP mainly depends on the density of markers used to calculate the GRMs in the structured population 17,31. To improve computing efficiency, several simplified algorithms such as FaST-LMM, GRAMMAR-Gamma and BOLT-LMM sampled or screened a small proportion of the whole genomic SNPs to estimate the GBVs or genomic heritability as precisely as possible. To underestimate the GBVs, GRAMMAR-Lambda would require fewer sampling markers than the competing methods, as demonstrated by simulation (3), wherein GRAMMAR-Lambda pre-specified a genomic heritability of 0.5.

We randomly sampled 3,000, 5,000, 10,000, 15,000, 20,000, and 25,000 SNPs from entire genomic markers to calculate the GRMs. Figure 3 shows the changes in the genomic controls of GRAMMAR-Lambda and four competing methods with numbers of sampling markers. No competing method could control false-positive/negative errors with less than 50,000 sampling SNPs. Specifically, both FaST-LMM and GRAMMAR-Gamma gradually controlled false-positive errors with the increasing number of sampling markers. GRAMMAR calibrated the false-negative rate by under-estimating GBVs with fewer markers, while BOLT-LMM yielded a high rate of false-positive errors due to inflating test statistics, independent of where the markers were drawn from. In comparison, GRAMMAR-Lambda retained reasonable powers to detect QTNs through prefect genomic control, even with fewer than 3,000 sampling markers (see Figures 4S and 5S).

**Figure 4:**
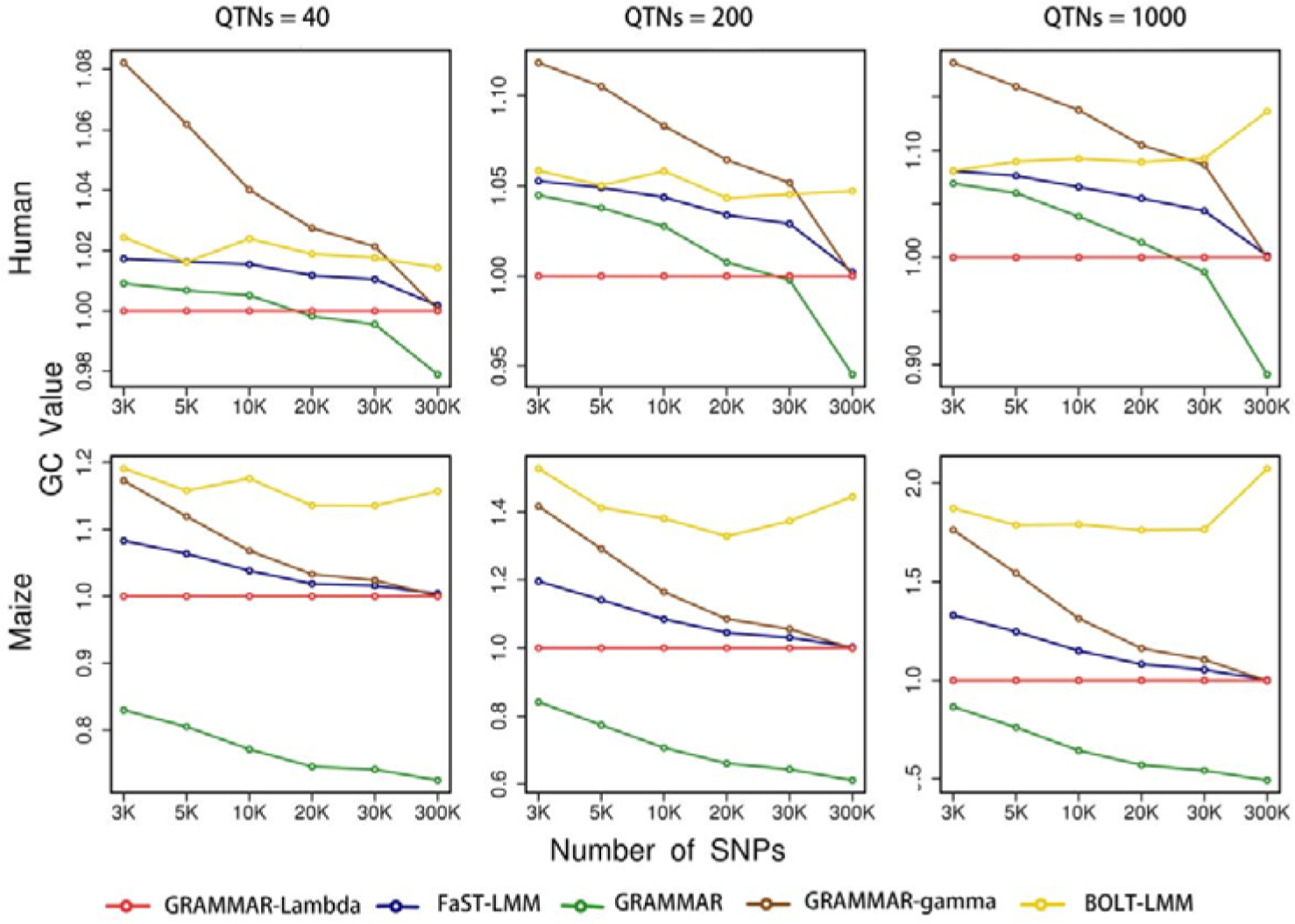
Changes in genomic controls with the number of sampling SNPs for GRAMMAR-Lambda and the four competing methods. Genomic control is calculated by averaging genome-wide test statistics. The simulated phenotypes are controlled by 40, 200 and 1,000 QTNs with the moderate heritability in human and maize.

### Application to real data

Previously published datasets of *Arabidopsis thaliana*, mice, and maize were used to illustrate GRAMMAR-Lambda and compare it with FaST-LMM, GRAMMAR, GRAMMAR-Gamma and BOLT-LMM in terms of QTN mapping and genomic control. Thirty-two phenotypes with greater than 120 records were selected for GWAS for *Arabidopsis thaliana* and 109 phenotypes for mice using the normality test. For the two datasets, we did not record computing times, because either a small population or a few markers were enough to make a distinction between these competing methods.

Using all compared methods, we depicted Q-Q and Manhattan profiles for the traits of detectable QTNs (Figures 6S–8S for *Arabidopsis thaliana*, mice, and maize, respectively). GRAMMAR, GRAMMAR-Gamma, and BOLT-LMM performed almost the same statistical properties as in simulations. GRAMMAR detected a handful QTNs with substantial false-negative errors, while BOLT-LMM detected more QTNs with higher false-positive errors than other competing methods but still fewer QTNs than GRAMMAR-Lambda with joint association analysis. Under reasonable genomic control, overall, GRAMMAR-Gamma identified fewer QTNs than GRAMMAR-Lambda with an association test at a time. In what follows, we focused on comparing the mapping results obtained with GRAMMAR-Lambda and exact FaST-LMM.

As shown in the Q-Q profiles, GRAMMAR-Lambda with either one test at a time or joint analysis performed almost perfect genomic control in the exact or infinitely close to the averaged Chi-squared statistics of 1.0. Using GRAMMAR-Lambda with joint association analyses, we found QTNs in 21 of 32 phenotypes of *Arabidopsis thaliana* and 56 of 109 phenotypes of mice. However, FaST-LMM did not find any QTN in 1 and 5 of these 21 and 56 traits, respectively. Meanwhile, GRAMMAR-Lambda identified 32 and 242, respectively, more QTNs with joint association analyses than exact FaST-LMM. Even with one association test at a time, GRAMMAR-Lambda covered 90% of the QTNs detected with FaST-LMM for mice phenotypes, while about 60% for *Arabidopsis thaliana* phenotypes. Notably, FaST-LMM produced unstable genomic controls for the analysed traits in *Arabidopsis thaliana*.

To analyse the flowering time in maize, we implemented GRAMMAR-Lambda in estimating genomic heritability and GBVs together by using entire genomic markers or estimating only GBVs by randomly sampling 5,000 SNPs with an empirical heritability of 0.8. GRAMMAR-Lambda, one association test at a time detected the same QTNs as GRAMMAR-Gamma and completely covered 4 QTNs distributed on chromosomes 3, 6, 8, 10 identified with exact FaST-LMM. The GRAMMAR-Lambda revealed fewer QTNs with sampling markers than those with entire genomic markers by a two-QTN difference, but joint association analysis could separate more signals corresponding to QTNs detected with either entire or sampling markers than a test at once. From the input of genotypes and phenotypes to the output of QTN mapping, the GWAS for flowering time consumed computing time of 2.073, 3.637, 0.101, and 0.058 min with GRAMMAR, GRAMMAR-Gamma, and the GRAMMAR-Lambdas by using entire and sampling markers. Additionally, FaST-LMM ran for 32.147 min in Single-RunKing ^32^ and BOLT-LMM for 166.448 min in BOLT ^27^.

## Discussion

The modified GRAMMARs, GRAMMAR-Gamma, and BOLT-LMM have the same computing complexity for association tests as GRAMMAR-Lambda, but both methods require estimated variance components or polygenic heritability. To prevent false-positive errors, GRAMMAR-Gamma is not used to calculate the GRM with fewer selected markers. Although BOLT-LMM estimates the calibration factor with only 30 pseudorandom SNPs in human datasets ^27^, it produces a high false-positive error rate in a complex structured population. Extensive simulations demonstrated the following: 1) In estimating GBVs, variance components or genomic heritability could be empirically specified over a broad range, without estimation in advance; 2) even using fewer randomly drawn markers from the whole genome to calculate the GRM, GRAMMAR-Lambda had almost the same statistical power as the exact mixed-model association analysis.

Generally, computing costs are attributed to building the GRM, estimating variance components or polygenic heritability, and computing association statistics in genome-wide mixed-model association analysis ^33^. To date, no simplified algorithm has been published that can comprehensively reduce these three computational charges, while simultaneously improving the statistical power to detect QTNs. For a genomic dataset containing *m* SNPs genotyped on *n* individuals, GRAMMAR-Lambda took only the computing time of *O*(*mn*^2^) to build the GRM and *O*(*mn*) for association tests. Instead of the GBLUP, we adopt ridge regression ^34^ to estimate the GBVs, reducing the computing time to build the GRM to 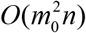, as in FaST-LMM-Select ^35^. When analysing a large-scale population with complex structure, we can simplify GBVs’ estimation in following way: with a specified heritability, to firstly solve the effects of *m*_0_ sampling markers 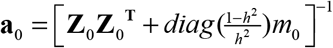 by using ridge regression ^34^, and then to calculate GBVs as **Z**_**0**_**a**_**0**_. As a result, such a negligible computational complexity to estimate the GBVs allows GRAMMAR-Lambda to implement large mixed model association analyses in naïve association tests of PLINK ^36^ or GCTA ^37^. A user-friendly GRAMMAR-Lambda software with these simplifications was developed, which is freely available at https://github.com/RunKingProgram/GRAMMAR-Lambda.

With the consideration of possible linkage disequilibrium among candidate markers, joint association analysis for multiple QTN candidates obtained with multiple testing significantly improves the statistical power to detect QTNs. Moreover, it is not required to re-estimate the residuals and genomic control in stepwise regression, generating a higher estimating robustness and computing efficiency than the Gaussian mixture-model association analysis ^38–40^ of BOLT-LMM. Since there is no need to estimate variance components (heritability or genetic relationship), GRAMMAR-Lambda can easily be extended to handle multiple correlated phenotypes and binary disease traits, which is expected to simplify genome-wide multivariate and generalized mixed-model association studies significantly.

## Acknowledgements

The research is financially supported by the National Key R&D Program of China (2018YFD0900201) and the National Natural Science Foundations of China (32072726).

## Online Methods

### Genomic mixed-model

Before genome-wide association analysis, the phenotype is generally justified for some fixed factors, such as ethnicity, sex and age. For comprehensive genomic control of population stratification, family structure and cryptic relatedness, we described the relationship between phenotype and the tested SNPs using the following LMM:

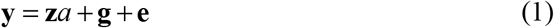

where **y** is a vector of the justified phenotype; a is a scalar for genetic effect of the tested SNPs on phenotype; **z** is a vector of indicator variables for SNP genotypes, which is generally coded as 0, 1 or 2 for the three genotypes AA, AB and BB; **g** is a vector of polygenic effects excluding the tested SNP; 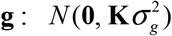 with 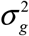 being the polygenic variance and **K** the GRM among individuals; **e** is the residual error distributed as 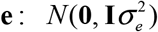 with 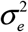 being the residual variance and **I** the identity matrix.

### GRAMMAR algorithm

In essence, GRAMMAR solves genomic mixed model using the two-stage regression method. The first stage is to estimate polygenic effects with the null model:

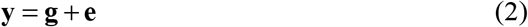

To avoid computing the inverse of GRM, we construct GBLUP equations as

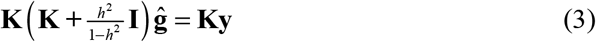

where *h*^2^ is the genomic heritability. Equation (3) can be solved using the gradient descent algorithm for a large population.

The second stage is to statistically infer association of resulting phenotypic residuals at the first stage with the tested SNP using simple regression model:

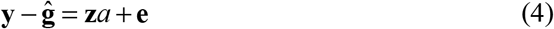

We formulate test statistic as

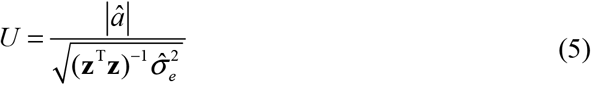

with *â* = (**z**^T^**z**)^−1^**z**^T^ (**y** − **ĝ**) and 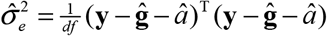, where *df* is residual degree of freedom.

The U statistic subjects to a standard normal distribution and in what of follows, we transform normal *U* to Chi-squared statistic *χ*^2^ = *U*^2^ to evaluate genomic control.

### Genomic control

At the first stage of GRAMMAR, we estimate the GBVs rather than polygenic effects. in addition of polygenic effects, as known, there are a part of QTN effects in the GBVs. Moving partial QTN effects from phenotypic residuals makes genome-wide test statistics to be deflated at the second stage, which yields serious false-negative error, especially when GEBV was precisely estimated for complex population structure. In this case, the test statistic *U* will subject to

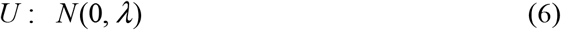

rather than the standard normal distribution, where *λ* is the variance deflation factor (so-called genomic control). According to the frequentist outlier test ^1^, we could efficiently control the deflation of the test statistics *χ*^2^ by

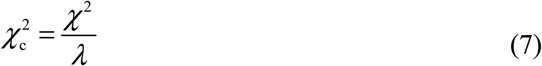

where *λ* is estimated as the mean or median of *χ*^2^ obtained with the GRAMMAR. 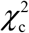 follows Chi-squared distributions with the degree of freedom 1.

### Joint association analysis

After genomic control for GRAMMAR, some SNPs with relatively low linkage disequilibrium were chosen as QTN candidates. In general, a lot of markers may be chosen as QTN candidates at the significance level lower than stringent Bonferroni corrected criterion ^2^, but number of QTN candidates should not be limited to less than population size to enhance efficiency for model selection. For the residual **y** − ĝobtained with the GBLUP, we jointly analysed multiple QTN candidates to improve the statistical power to detect QTNs. Backward regression was adopted to optimise the following multiple linear model:

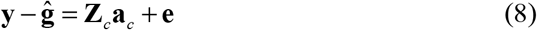

where, **Z**_*c*_**a**_*c*_ is the regression terms of QTN candidates. Given the Bonferroni corrected level of significance, the genetic effects were selected stepwise, and corresponding QTNs identified according to the corrected statistics (7).

## Simulations

To verify the adaptability of GRAMMAR-Lambda to population structure, we used human genomic data ^3^ with simple population stratification, and maize genomic data ^4^ with complex cryptic relativeness among individuals to simulate trait phenotypes. With higher quality control, 300,000 SNPs were extracted from both 3000 human and 2640 maize. QTNs were randomly distributed over the whole genomic SNPs. Additive effects of simulated QTNs were drawn from the gamma distribution with shape=1.66 and scale=0.4 so that there were few with large effects and more with minor effects in the simulated QTNs. The genetic effect for each individual was calculated as the sum of all QTN genotypic effects. In sampling residual errors using the normal distribution, the residual variance was regulated by the given genomic heritability of traits. The trait phenotypes were generated by adding up the genetic effects and residual errors.

In addition to population structure, the number of QTNs, genomic heritability and sampling number of SNPs were considered as experimental factors in the simulations. We carried out three simulations to investigate (1) statistical properties of GRAMMAR-Lambda by simulating phenotypes controlled by 40, 200 and 1000 QTNs with varied levels of heritabilities (low (0.2), moderate (0.5), and high (0.8)); (2) sensitivity to estimates of variance components or genomic heritability by using the phenotypes controlled by 200 QTNs in the first simulation; and (3) calculation of GRMs with sampling markers by using the phenotypes at the moderate genomic heritability in the first simulation. In the first and third simulations, GRAMMAR-Lambda was compared to FaST-LMM, GRAMMAR, GRAMMAR-Gamma and BOLT-LMM in terms of Q-Q profiles, or genomic controls, and ROC profiles. Especially in the first simulation, GRAMMAR-Gamma jointly analyses multiple QTN candidates with a test at once.

Under perfect genomic control (1.0), the ROC profiles can be plotted using the statistical powers to detect QTNs relative to the given series of Type I errors. Statistical powers were defined as the percentage of identified QTNs that had the maximum test statistics among their 20 closest neighbours over the total number of simulated QTNs. Simulations were repeated 50 times, in each the positions and effects of simulated QTNs varied, and the average results were recorded.

## Real data

The datasets were collected from three species: 1) the *Arabidopsis thaliana* dataset included 199 samples, with 216,130 SNPs and 107 phenotypes ^5^; 2) the mouse dataset had 1,940 samples with 12,226 SNPs and 123 phenotypes ^6^; 3) the maize datasets involved 2,279 inbred lines, each with 681,258 SNPs genotyped and the flowering time phenotype measured as days to silk ^4^ (URL: http://www.panzea.org/!#genotypes/cctl). All data analyses were performed in a CentOS Linux sever with 2.60 GHz Intel(R) Xeon(R) 40 CPUs E5-2660 v3, and 512 GB.

